# Weighted burden analysis of exome-sequenced case-control sample implicates synaptic genes in schizophrenia aetiology

**DOI:** 10.1101/203521

**Authors:** David Curtis, Leda Coelewij, Shou-Hwa Liu, Jack Humphrey, Richard Mott

## Abstract

A previous study of exome-sequenced schizophrenia cases and controls reported an excess of singleton, gene-disruptive variants among cases, concentrated in particular gene sets. The dataset included a number of subjects with a substantial Finnish contribution to ancestry. We have reanalysed the same dataset after removal of these subjects and we have also included non-singleton variants of all types using a weighted burden test which assigns higher weights to variants predicted to have a greater effect on protein function. We investigated the same 31 gene sets as previously and also 1454 GO gene sets. The reduced dataset consisted of 4225 cases and 5834 controls. No individual variants or genes were significantly enriched in cases but 13 out of the 31 gene sets were significant after Bonferroni correction and the “FMRP targets” set produced a signed log p value (SLP) of 7.1. The gene within this set with the highest SLP, equal to 3.4, was *FYN*, which codes for a tyrosine kinase which phosphorylates glutamate metabotropic receptors and ionotropic NMDA receptors, thus modulating their trafficking, subcellular distribution and function. In the most recent GWAS of schizophrenia it was identified as a “prioritized candidate gene”. Two of the subunits of the NMDA receptor which are substrates of *FYN* are coded for by *GRIN1* (SLP=1.7) and *GRIN2B* (SLP=2.1). Of note, for some sets there was a substantial enrichment of non-singleton variants. Of 1454 GO gene sets, 3 were significant after Bonferroni correction. Identifying specific genes and variants will depend on genotyping them in larger samples and/or demonstrating that they cosegregate with illness within pedigrees.

## Introduction

Schizophrenia is a severe and disabling mental illness with onset typically in early adult life. It is associated with low fecundity but nevertheless remains fairly common with a lifetime prevalence of around 1% (Power et al., 2013). A variety of types of genetic variation contribute to risk. Many common variants demonstrate association with small effect sizes whereas extremely rare variants can have very large effect sizes. 108 SNPs have been reported to be genome-wide significant with odds ratio (OR) of 1.1-1.2 and it is likely that many other variants will achieve statistical significance when larger samples are genotyped (Schizophrenia Working Group of the Psychiatric Genomics Consortium, 2014). Weak effects from common variants may arise from a number of mechanisms. The variant itself may exert a direct effect at some point in the pathogenic process, it may pick a up a more indirect effect through involvement in gene regulatory networks or it may be in linkage disequilibrium with other variants have a larger, direct effect (Boyle et al., 2017). A recent example of the last case is provided by SNPs in the HLA region which tag variant haplotypes of C4, the gene for complement component 4, the different haplotypes producing different levels of C4A expression associated with OR for schizophrenia risk of 1.3 (Sekar et al., 2016). Variants associated with small effect sizes will be subject to relatively little selection pressure and hence can remain common. By contrast, extremely rare variants such as some copy number variants (CNVs) or loss of function (LOF) variants of SETD1A may lead to a very high risk of developing schizophrenia (Deciphering Developmental Disorders Study, 2017; Rees et al., 2014; Singh et al., 2016). A proportion of cases of schizophrenia seem to be due to such variants with large effect size arising as de novo mutations (DNMs) (Fromer et al., 2014; Singh et al., 2017). Such variants are likely to be subject to strong selection pressure and may only persist for a small number of generations. Theoretically, variants acting recessively might persist in the population and still have reasonably large effect size but attempts to identify these have to date been unsuccessful (Curtis, 2015; Rees et al., 2015; Ruderfer et al., 2014).

In order to focus attention on only new or recent variants, the Swedish schizophrenia study of whole exome sequence data focussed on what were termed ultra-rare variants (URVs), that is variants which only occurred in a single subject and which were absent from ExAC. The effects of some of these variants on gene function were annotated as damaging or disruptive and these variants, termed dURVs, were found to be commoner in cases than controls across all genes, with the effect concentrated in particular sets of genes including FMRP targets, synaptically localised genes and genes which were LOF intolerant (Genovese et al., 2016). The present study seeks to analyse this dataset further in order to consider whether rare non-singleton sequence variants, as well as singleton variants, contribute to schizophrenia risk.

The dataset used in this study overlaps with a number of previously reported analyses. The full exome-sequenced dataset consists of 4968 cases with schizophrenia and 6245 controls. Although recruited in Sweden, it should be noted that some subjects have a substantial Finnish component to their ancestry (Genovese et al., 2016). The earlier phase of this dataset consisted of 2045 cases and 2045 controls and the primary analysis of these revealed subjects an excess among cases of very rare, disruptive mutations spread over a number of different genes though concentrated in particular gene sets (Purcell et al., 2014). This first phase of the dataset was also used for analyses which attempted to detect recessive effects and to identify Gene Ontology (GO) pathways with an excess of rare, functional variants among cases but which did not produce statistically significant results (Curtis, 2016, 2013). A subset of the full dataset with cases with Finnish ancestry removed was used to demonstrate a method for deriving an exome-wide risk score and to demonstrate an association of schizophrenia with variants in mir134 binding sites (Curtis, 2017; Curtis and Emmett, 2017). A genetically homogeneous subset of the full Swedish dataset was combined with a UK case-control association sample and nonsynonymous variants with Minor Allele Frequency (MAF)<0.001 which were present on the Illumina HumanExome and HumanOmniExpressExome arrays were analysed (Leonenko et al., 2017). This revealed an enrichment of these variant alleles in LOF intolerant genes and FMRP targets.

The present study uses a subset of the Swedish dataset after removal of subjects with a high Finnish ancestry component in order to avoid artefactual results produced by population stratification. It also utilises all rare (MAF<0.01) variants analysed using a weighted burden test to identify genes and sets of genes associated with schizophrenia risk.

## Methods

The data analysed consisted of whole exome sequence variants downloaded from dbGaP from the Swedish schizophrenia association study containing 4968 cases and 6245 controls (Genovese et al., 2016). The dataset was managed and annotated using the GENEVARASSOC program which accompanies SCOREASSOC (https://github.com/davenomiddlenamecurtis/geneVarAssoc). Version hg19 of the reference human genome sequence and RefSeq genes were used to select variants on a gene-wise basis. Members of the protocadherin gamma gene cluster, whose transcripts overlap each other but which are entered separately in RefSeq, were treated as a single gene which was labelled PCDHG.

A number of QC processes were applied. Variants were excluded if they did not have a PASS in the Variant Call Format (VCF) information field and individual genotype calls were excluded if they had a quality score less than 30. Sites were also excluded if there were more than 10% of genotypes missing or of low quality in either cases or controls or if the heterozygote count was smaller than both homozygote counts in both cohorts. As previously reported (Curtis, 2017), preliminary gene-wise weighted burden tests revealed that several genes had an apparent excess of rare, protein-altering variants in cases but that these results were driven by variants which were reported in ExAC to be commoner in Finnish as opposed to non-Finnish Europeans (Lek et al., 2016). Accordingly, subjects with an excess of alleles more frequent in Finns were identified and removed from the dataset, comprising 743 cases and 411 controls. Once this had been done, leaving a sample of 4225 cases and 5834 controls, the gene-wise weighted burden test results conformed well to what would be expected under the null hypothesis with no evidence for inflation of the test statistic across the majority of genes not thought to be implicated in disease.

The tests previously carried out for an excess of dURVs among cases (Genovese et al., 2016) were performed on both the full and reduced datasets, with and without including covariates consisting of the total URV count and the first 20 principal components from the SNP and indel genotypes.

Weighted burden analysis of genes and gene sets as described below was carried out using SCOREASSOC, which analyses all variants simultaneously and can accord each variant a different weight according to its MAF and its predicted function (Curtis, 2016, 2012). Each variant was annotated using VEP, PolyPhen and SIFT (Adzhubei et al., 2013; Kumar et al., 2009; McLaren et al., 2016). GENEVARASSOC was used to generate the input files for SCOREASSOC and the default weights were used, for example consisting of 5 for a synonymous variant and 20 for a stop gained variant, except that 10 was added to the weight if the PolyPhen annotation was possibly or probably damaging and also if the SIFT annotation was deleterious. The full set of weights is shown in supplementary Table S1. SCOREASSOC also weights variants more highly than common ones but because it is well-established that no common variants have a large effect on the risk of schizophrenia we excluded variants with MAF>0.01 in the cases and in the controls, so in practice weighting by rarity had negligible effect. For each subject a gene-wise risk score was derived as the sum of the variant-wise weights, each multiplied by the number of alleles of the variant which the given subject possessed. These scores were then compared between cases and controls using a t test. To indicate the strength of evidence in favour of an excess of rare, functional variants in cases we took the logarithm base 10 of the p value from this t test and then gave it a positive sign if the average weighted sum was higher in cases and a negative sign if the average was higher in controls, to produce a signed log p (SLP).

In order to explore the contribution of singleton variants, for the analyses of gene sets three sets of variants were used: singleton variants which were only observed in a single subject and not in ExAC; non-singleton variants, observed in more than one subject (though still with MAF<0.01 in cases and/or controls); all variants, consisting of these singleton and non-singleton combined.

Weighted burden analysis within sets of genes was carried out using PATHWAYASSOC, which for each subject sums up the gene-wise scores to produce an overall score for the gene set. These set-wise scores can then be compared between cases and controls using a t-test. This approach has been demonstrated to produce appropriate p values through application to real data, supported by permutation testing (Curtis, 2016). This analysis was applied to the 31 gene sets used in the Swedish study separately using singleton, non-singleton and all variants. The analysis was also applied using all variants to the 1454 "all GO gene sets, gene symbols" pathways downloaded from the Molecular Signatures Database at http://www.broadinstitute.org/gsea/msigdb/collections.jsp (Subramanian, Tamayo et al. 2005).

Logistic regression analyses of dURVs were carried out using R (R Core Team, 2014). Weighted burden tests for genes and gene sets were carried out using SCOREASSOC and PATHWAYASSOC. Results from these programs are expressed as a Signed Log P (SLP) which is positive if there is an excess of variants among cases and negative if there is an excess among controls. Thus, a SLP of 3 would indicate that there was an excess of variants among cases with two-tailed significance P<10^-3^.

## Results

Preliminary analysis of the whole dataset, (i.e. all individuals before excluding those with Finnish ancestry), using a logistic regression analysis to test for an excess of dURVs among cases was significant (P=8.7*10^-10^) when the total URV count and principal components were included as covariates. However without covariates this analysis was only marginally significant (P=0.031). Further investigation showed that subjects with a substantial Finnish component to their ancestry had a larger number of URVs than those who did not. Cases tended to have a larger number of dURVs than controls, but only relative to the total number of URVs, and more cases had a substantial Finnish ancestry component than controls. Thus, in the whole sample the relative excess of dURVs among cases was almost completely masked by the fact that more cases had Finnish ancestry and that these cases had a smaller absolute number of URVs, meaning that overall there was only a small excess in the absolute number of dURVs among cases. Including the total URV count or the principal components or both as covariates allowed the relative excess among cases to become apparent. The analysis was then repeated on the reduced dataset without those subjects with a substantial Finnish ancestry component. Once this had been done, there was a significant absolute excess of dURVs among cases (P=2.7*10^-5^), without needing to include either total URV count or principal components as covariates.

The weighted burden tests evaluated 1,042,483 valid variants in 22,023 genes. As described in the *Methods* section, in preliminary analyses using the full dataset a number of genes yielded high SLPs. An example was *COMT*, with SLP=7.4. On inspection, it seemed that this gene-wise result was largely driven by SNP rs6267, which was heterozygous in 51/6242 controls and 94/4962 cases (OR=2.3, p=8*10^-7^). However this variant is noted in ExAC to have MAF=0.002 in non-Finnish Europeans but MAF=0.05 in Finns. Hence, its increased frequency among cases appeared to be due to the excess of cases with Finnish ancestry. Once all subjects with a substantial Finnish ancestry component were excluded, the SLP for *COMT* fell to 1.7 and for rs6267 there were 36/5831 heterozygous controls s and 36/4221 cases (OR=1.4, p=0.2). A similar effect was observed for other genes with excessively high SLPs in the full dataset but not in the reduced dataset, suggesting that removing subjects with substantial Finnish ancestry seemed to produce a satisfactorily homogeneous dataset. QQ-plots for the gene-wise analyses using the reduced dataset are shown in Figure 1. All of the plots are symmetrical, indicating that the test is unbiased. When only singleton variants are used the gene-wise tests are somewhat underpowered and the gradient is less than 1. However for the tests using non-singleton variants or all variants the SLPs almost exactly follow the distribution expected under the null hypothesis. One outlier is apparent. This is caused by the gene *CDCA8* which produces an SLP of -5.49 with all variants. Further inspection showed that this result was mainly driven by 22 highly weighted variant alleles among controls but only 5 among cases. For a gene-wise test to be exome-wide significant with 22,023 genes the absolute value of the SLP would need to exceed 5.64, so this result is still within chance expectation.

**Figure 1.**
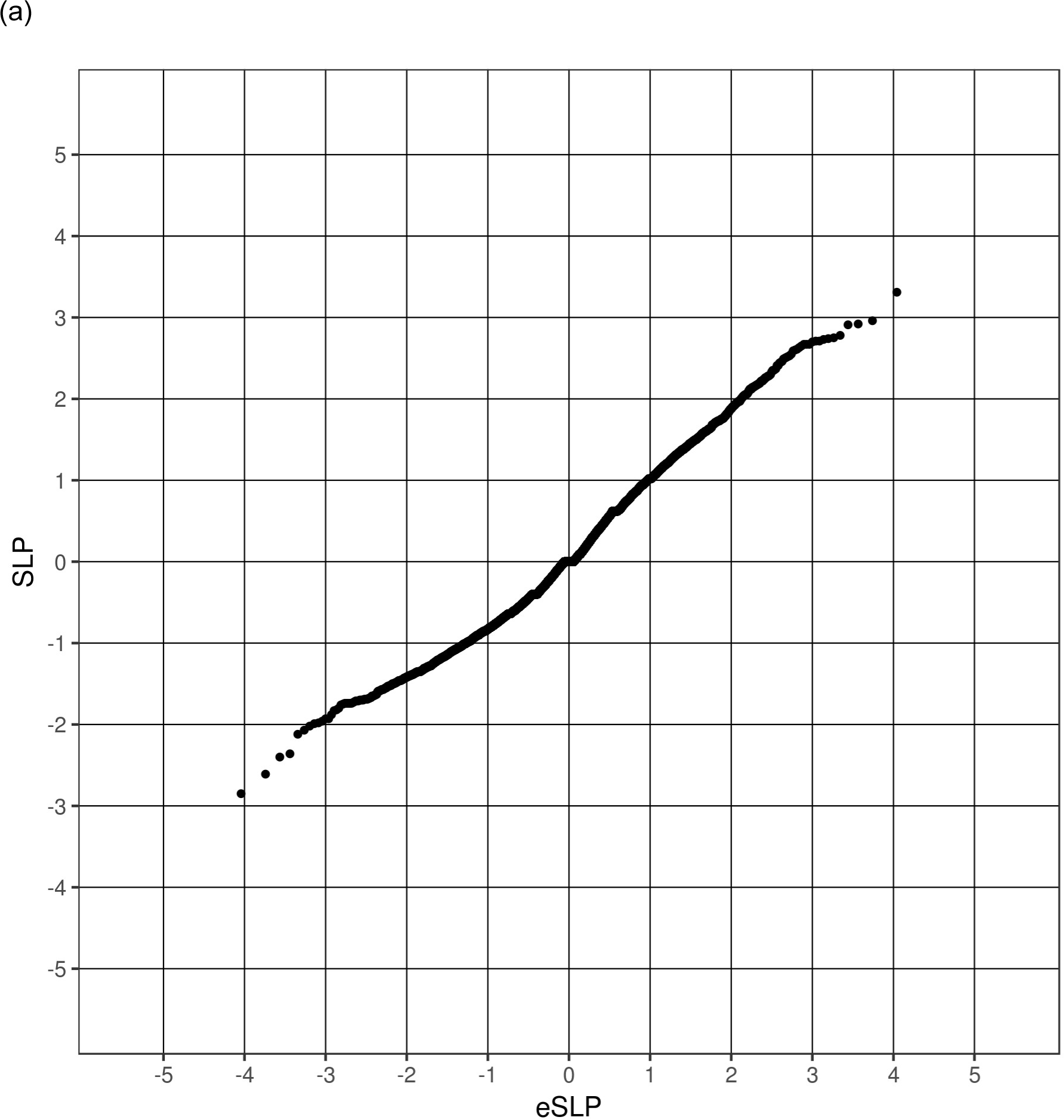
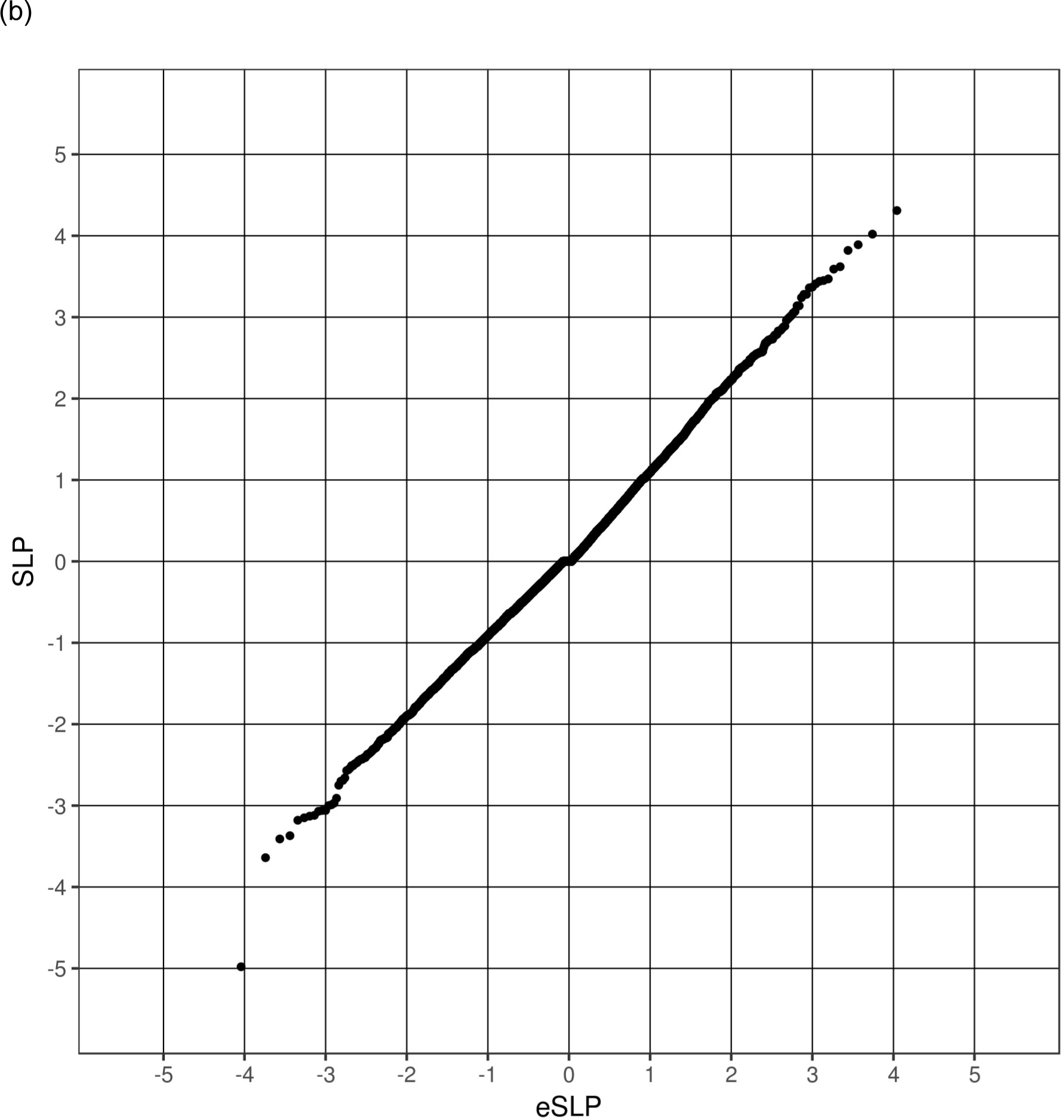
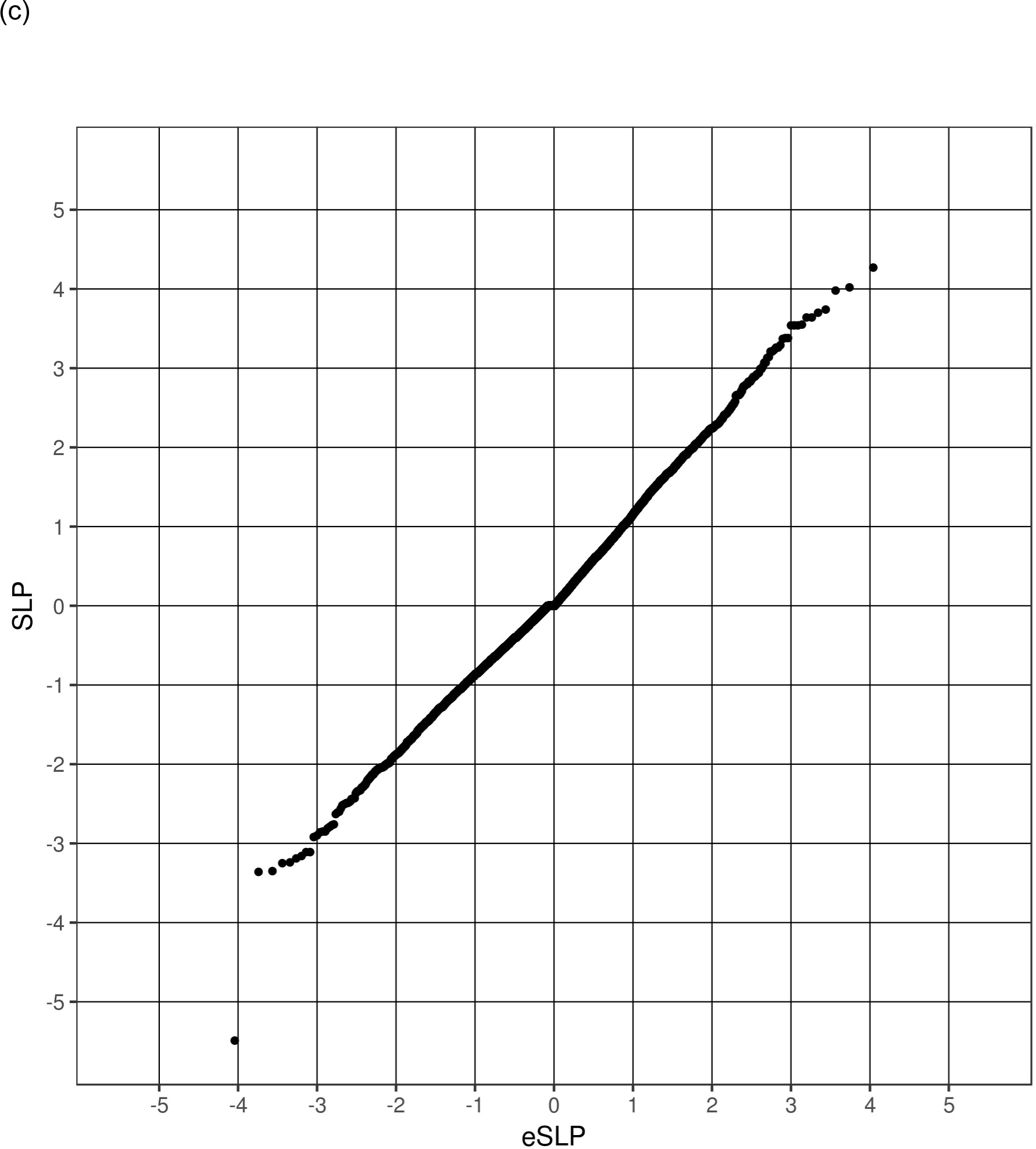
QQ plots of observed versus expected gene-wise SLP using (a) only singleton variants, (b) non-singleton variants and (c) both.

The results for the 31 gene sets which had previously been used in the Swedish study are shown in Table 1. Using the weighted burden test many, though not all, of the sets show an excess of variants among cases. For *neurons*, *pLI09*, *fmrp* and *mir137* the non-singleton variants make a substantial contribution but for *psd*, *rbfox13* and *rbfox2* the bulk of the effect comes from only the singleton variants. Given that there are 31 sets, a simple Bonferroni correction would mean that a set could be declared statistically significant if the SLP using all variants exceeded -log(31/0.05)=2.8 although this threshold should be regarded as conservative because the sets overlap each other. For the 13 sets where SLP>2.8 using all variants, the genes with the highest gene-wise SLPs are shown in Table 2. As expected, there is some overlap between the sets with several genes making contributions to more than one set. The gene with the highest gene-wise SLP in the *fmrp* set is *FYN* (SLP=3.4) and it is also a member of 6 other sets. *FYN* codes for a tyrosine kinase which phosphorylates glutamate metabotropic receptors and ionotropic NMDA receptors, which modulates their trafficking, subcellular distribution and function (Mao and Wang, 2016a) In the most recent GWAS of schizophrenia *FYN* was identified as a "prioritized candidate gene" and an intronic marker, rs7757969, was significant at p=4.8*10^-8^ (Li et al., 2017). The activity of *FYN* is regulated by dopamine DRD2 receptors (Mao and Wang, 2016b). *FYN* is involved in neuronal apoptosis, brain development and synaptic transmission and lower expression has been observed in the platelets of schizophrenic patients compared with controls (Ali and Salter, 2001; Du et al., 2012; Hattori et al., 2009). Two of the subunits of the NMDA receptor which are substrates of *FYN* are coded for by *GRIN1* (SLP=1.7) and *GRIN2B* (SLP=2.1). In all three of these genes, the signal seems to be produced from a number of highly weighted variants which are individually commoner in cases but all are very rare, with MAF<0.001 even among cases, so it is not possible to identify any obvious candidate variants.

**Table 1.**
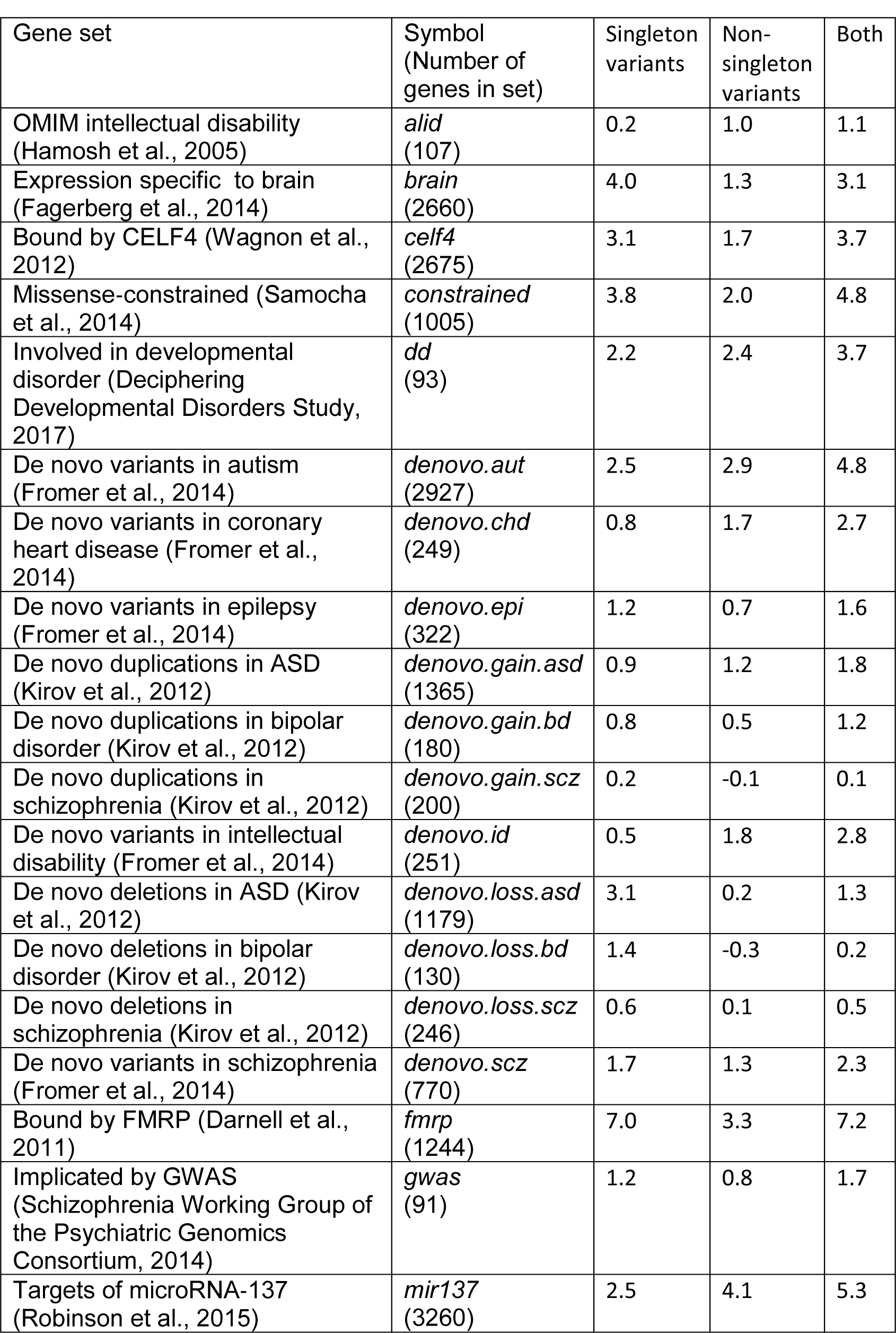
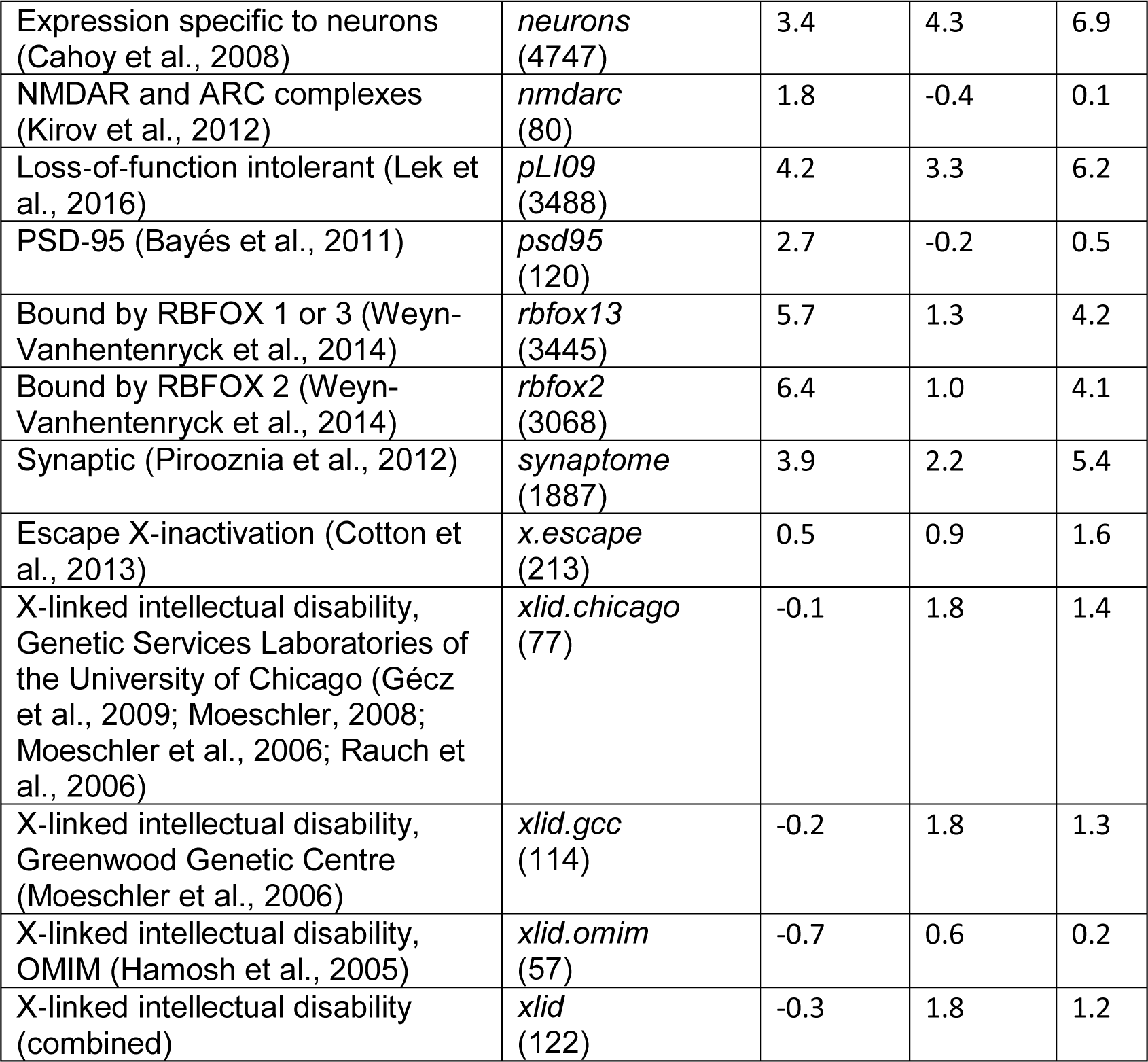
Results showing SLPs obtained for the gene sets used in in the original analysis Swedish schizophrenia study (Genovese et al., 2016). The lists of genes were obtained directly from the first author. The symbol used is the same as that used for the name of the file containing the list.

**Table 2.**
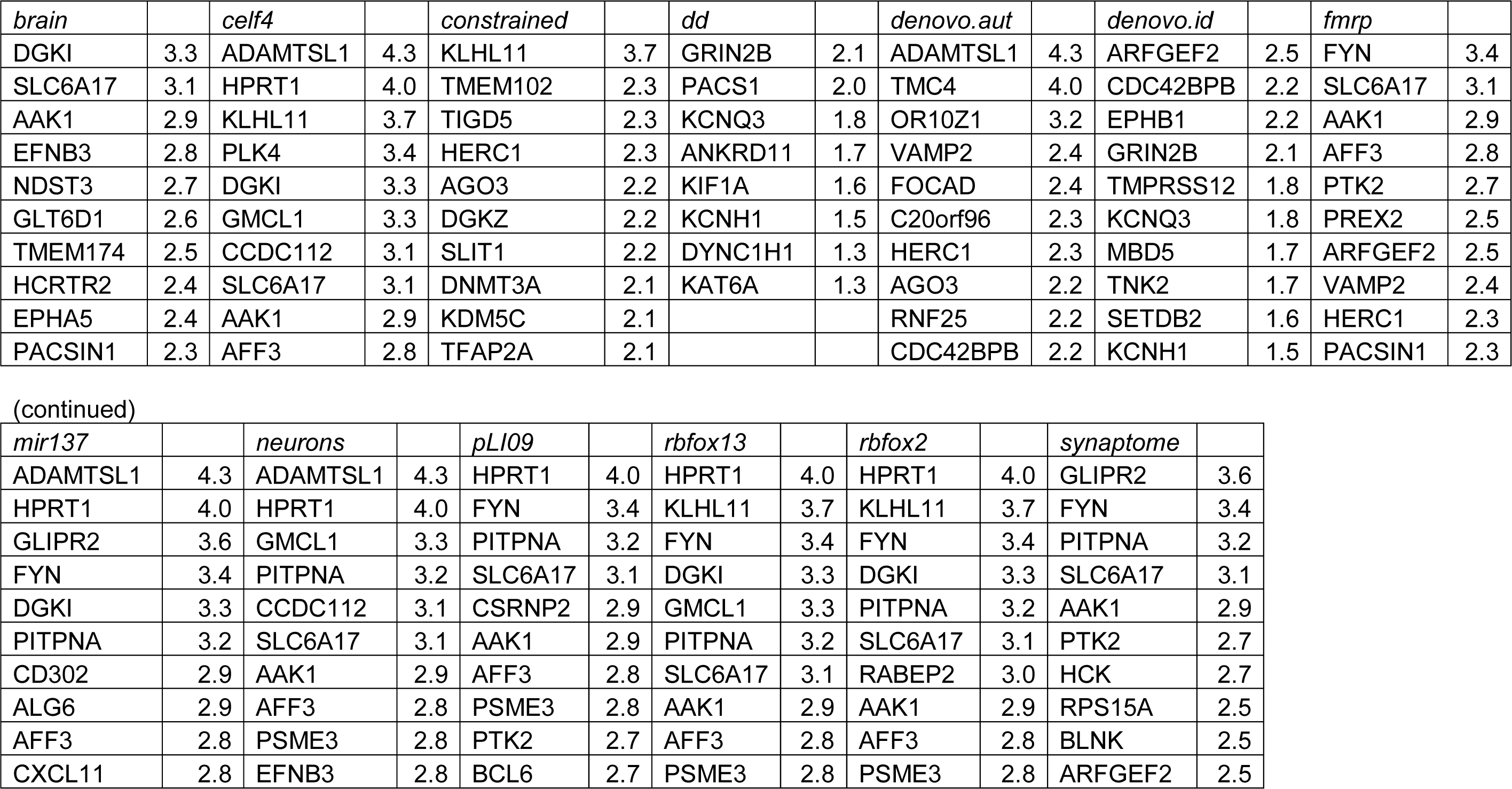
Gene-wise results for the genes with highest gene-wise SLPs in all sets with set-wise SLP>2.8. The top ten genes are shown, providing that the gene-wise SLP was at least 1.3, equivalent to p<0.05.

Figure 2 shows the QQ plot for the set-wise analyses using the GO gene sets. Given that there is overlap of genes between sets, the SLPs are non-independent and it is expected that the gradient of the QQ plot will be less than 1. For those sets with a negative SLP this is indeed the case and these results are in accordance with the expectation under the null hypoethesis. However the gradient becomes steeper for sets with positive SLPs and this can be interpreted as showing that some sets have an excess of variants among cases above that which would be expected by chance. Given that 1454 GO gene sets were tested, a simple Bonferroni correction would mean that a test could be declared "exome-wide significant" if it achieved an SLP exceeding -log(1454/0.05)=4.5. Three sets did achieve this threshold. However, given the fact that the set-wise SLPs are not independent a Bonferroni correction might be viewed as conservative and Table 3 shows all sets achieving SLP>3. The full results are presented in supplementary Table S2. The most significant set, INTRACELLULAR_SIGNALING_CASCADE with SLP=5.4, contains *FYN* and two other genes with gene-wise SLP>3, *S1PR4* (SLP=3.7) and *RTKN* (SLP=3.2). S1PR4 codes for the type 4 receptor for sphingosine-1-phosphate and the mouse strain carrying the mutation with genotype S1pr4^tm1Dgen^/S1pr4^+^ has decreased prepulse inhibition as a phenotype (http://www.informatics.jax.org/allele/genoview/MGI:3606610) (Blake et al., 2017; The Jackson Laboratory, n.d.). *RTKN* codes for rhotekin, a scaffold protein that interacts with GTP-bound Rho proteins. Again, inspecting results for individual variants within these genes did not reveal any obvious candidates. The full results for all genes and all gene sets can be downloaded at: http://www.davecurtis.net/downloads/SSS2WeightedBurdenAnalysisResults.zip.

**Figure 2.**
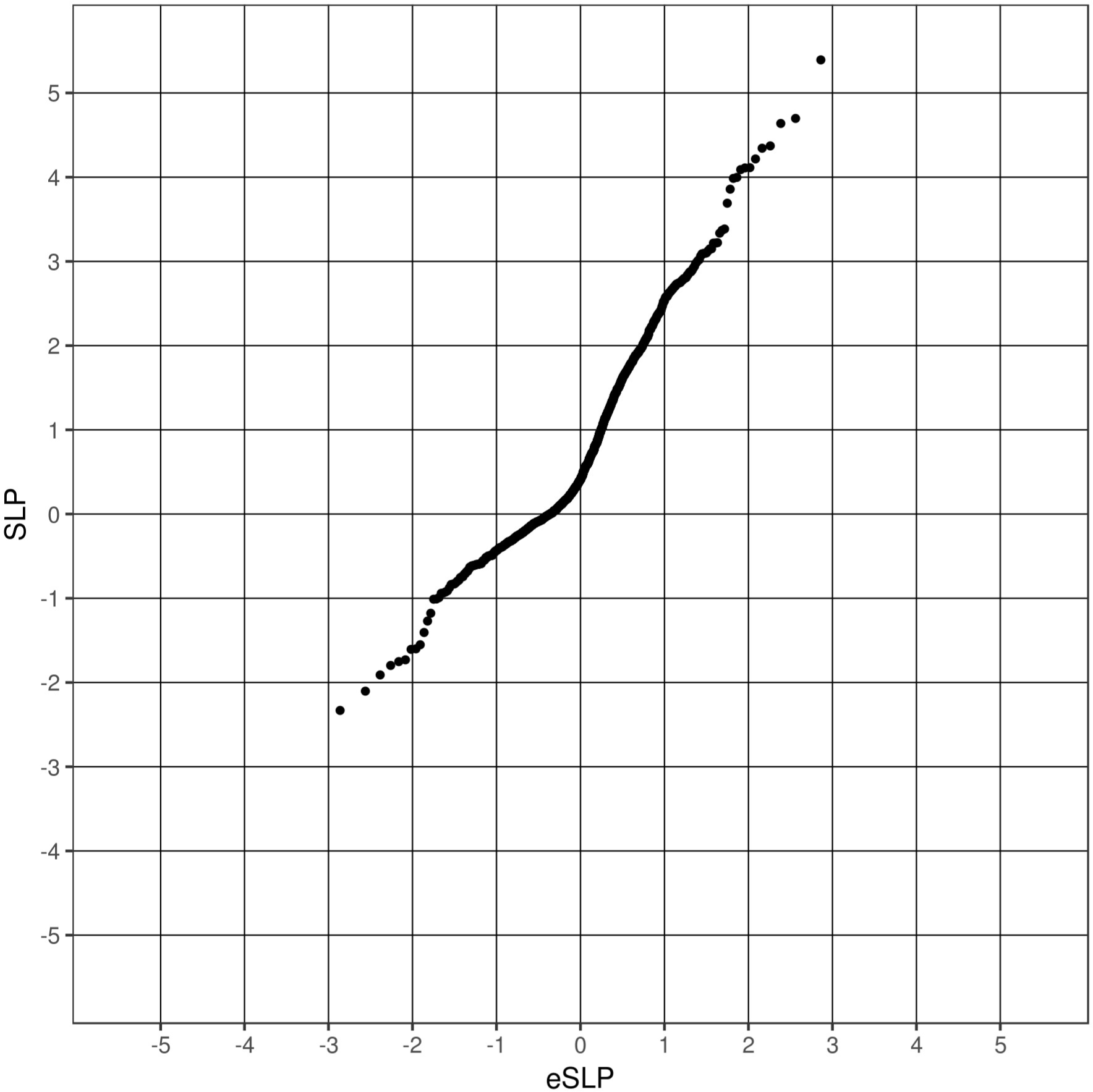
QQ plot for set-wise SLPs for GO sets against the expected SLP if all sets were non-overlapping and independent.

**Table 3.**
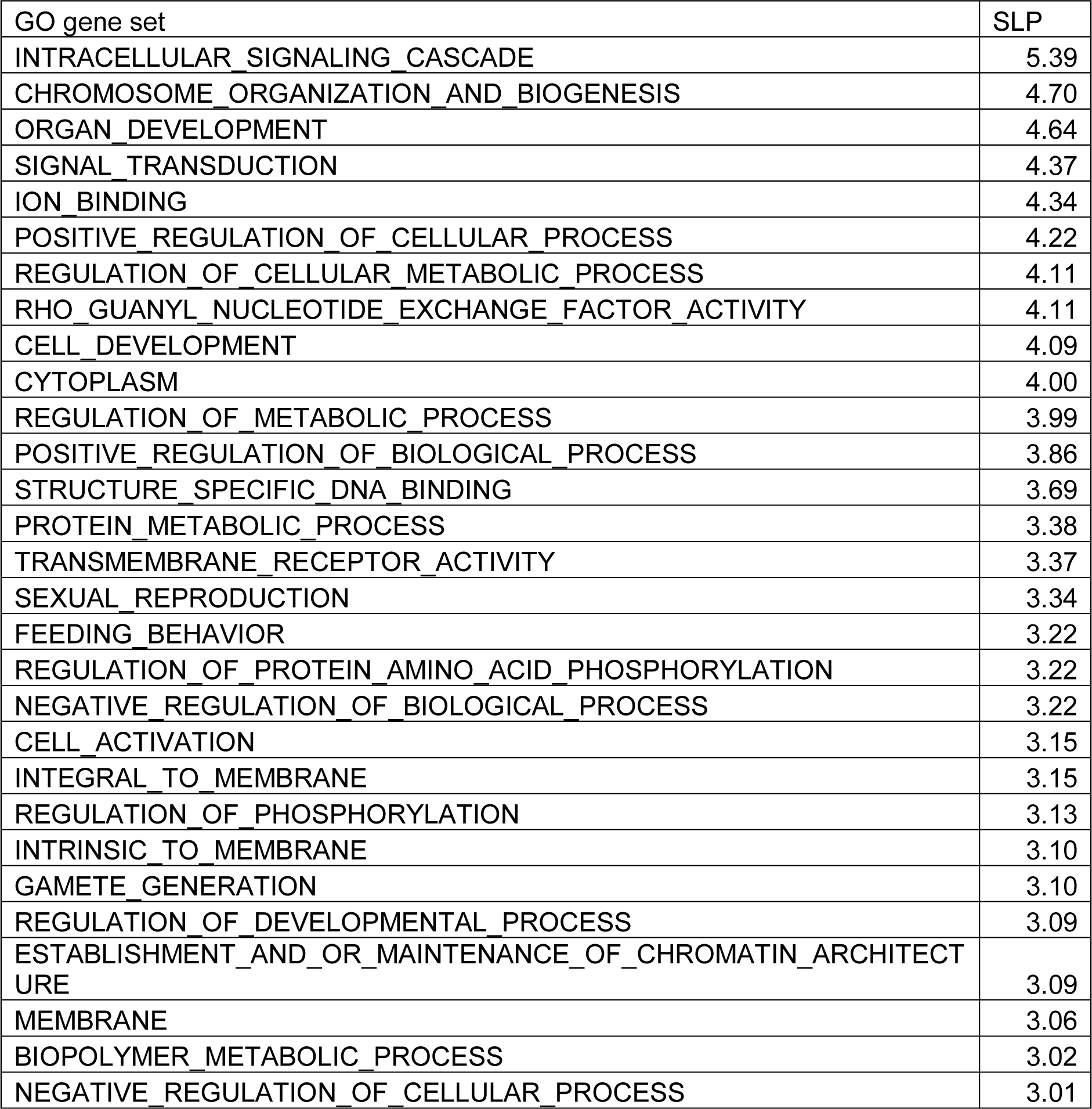
Table showing all of the 1454 GO gene sets which produced set-wise SLP>3.

## Discussion

This analysis identifies a number of sets of genes that meet Bonferroni-corrected criteria for statistical significance. It differs from previous analyses in a number of ways.

In contrast to the original analysis of the Swedish dataset (Genovese et al., 2016) it uses non-singleton as well as singleton variants and it clearly demonstrates that there is a contribution to risk from these non-singleton variants. This is extremely important in terms of the prospects for identifying rare risk variants for schizophrenia. If only unique variants conferred risk, that is only variants which occur independently as *de novo* mutations and then disappear after a small number of generations, then it would not be possible to identify any single variant as definitively affecting risk. One could at best identify perhaps classes of variant occurring in particular genes. Without being able to conclude that any particular variant affected risk, one could not carry out functional studies in model systems with the confidence that one was indeed studying a true risk variant. Additionally, if only unique variants contributed to risk then strategies that might use linkage disequilibrium to implicate untyped variants could not succeed. If, on the other hand, there are risk variants which survive and spread in the population then potentially these could be tagged by haplotypes of common SNPs and imputed from GWAS data, in a way similar to that used to impute C4 risk variants (Sekar et al., 2016). Alternatively population sequencing may soon become cheap and accurate enough to identify these rare variants directly.

This study differs from both the Swedish study (Genovese et al., 2016) and the Swedish-UK study (Leonenko et al., 2017) in that it uses a homogeneous dataset. The original study did not exclude the subjects with a substantial Finnish ancestry component whereas the Swedish-UK study did use a homogeneous subset of the Swedish subjects but then combined them with a UK sample. This meant that both studies needed to incorporate principal components to control for population stratification and this to some extent complicates the interpretation of their results. For example, the highly significant enrichment for dURVs reported in the first study only becomes apparent when covariates are included. In the Swedish-UK study, the most highly significant variant (p=3.4*10^-7^), which occurs in the *MCPH* gene, has MAF of 0.0046 in cases and of 0.0012 in controls, meaning that the unadjusted risk ratio is approximately 3.8. However after multivariate analysis including covariates the OR is reported as being only 1.2. By contrast, the reduced dataset we have used appears to be sufficiently homogeneous that the test statistic performs as expected without requiring any adjustment for population stratification. This allows for a simple, straightforward interpretation of the results obtained.

Another way our analysis differs is that it includes all variants in a single analysis. Variants are assigned different weights according to an arbitrary pre-specified set of weights designed to emphasise those variants more likely to affect gene function. This meant that we carried out only a single analysis for each gene or set of genes, reducing any correction for multiple-testing. Our analyses utilised 1,042,483 variants, compared with the 112,950 used in the Swedish-UK study. Using our method, 14 of the 32 candidate gene sets and 3 of the 1454 GO sets meet formal standards for statistical significance using a conservative Bonferroni correction.

As in the other studies, none of the results for individual genes reach formal standards for statistical significance, although the results obtained for *FYN* are possibly of interest. It seems likely that our results are detecting a real signal originating from rare variants concentrated within some of the genes that are members of the gene sets with high SLPs. These sets overlap each other to a considerable extent and it is difficult to tease out which ones best define a group of schizophrenia risk genes. An attempt to do this formally using exome-wide risk scores did not produce definitive results (Curtis, 2017). It should be noted that different sets might be implicated for different reasons. For example, it may be that the high SLP for targets of *miR-137* occurs because disruption of the regulation of these genes by *miR-137* can lead to increased risk of schizophrenia, as supported by the association of schizophrenia with markers for *miR-137* and with variants in its binding sites (Curtis and Emmett, 2017; Olde Loohuis et al., 2017). On the other hand, there is no reported association of *FMRP* itself with schizophrenia and the high SLP for its targets may simply reflect that this identifies a group of genes whose mRNA is localised to the synapse. In any event, it is clear that with samples currently available we are only able to identify very broad gene sets but not yet specific genes.

With increased sample sizes it will become possible to identify specific genes and variants which have a moderate or large effect on risk. However such variants, although not singletons, will still be very rare and serious attention should be focussed on complementary approaches to confirm them. One such approach would be to use exome sequence data from affected subjects to provide reference haplotypes for imputation into large GWAS datasets, analogously to the way C4 variants implicating risk were identified (Sekar et al., 2016). Another would be to search for affected relatives of subjects with candidate variants in order to see if the variants cosegregate with disease, a strategy which was successful in implicating *RBM12* in the aetiology of psychosis (Curtis, 2011; Steinberg et al., 2017). If and when specific variants are identified as having substantial effects on risk then they can be incorporated into model systems in order to gain insight into the mechanisms affecting the development of schizophrenia.

## Acknowledgements

The authors wish to thank Giulio Genovese for his assistance in providing supporting files and responding to queries. The datasets used for the analysis described in this manuscript were obtained from dbGaP at http://www.ncbi.nlm.nih.gov/gap through dbGaP accession number *phs000473.v2.p2*. Samples used for data analysis were provided by the Swedish Cohort Collection supported by the NIMH grant R01MH077139, the Sylvan C. Herman Foundation, the Stanley Medical Research Institute and The Swedish Research Council (grants 2009-4959 and 2011-4659). Support for the exome sequencing was provided by the NIMH Grand Opportunity grant RCMH089905, the Sylvan C. Herman Foundation, a grant from the Stanley Medical Research Institute and multiple gifts to the Stanley Center for Psychiatric Research at the Broad Institute of MIT and Harvard. JH is supported by MRC studentship 516702.

## Compliance with ethical standards

JH is supported by MRC studentship 516702. All procedures performed in studies involving human participants were in accordance with the ethical standards of the institutional and/or national research committee and with the 1964 Helsinki declaration and its later amendments or comparable ethical standards. Informed consent was obtained from all individual participants included in the study.

